# A workflow for accurate metabarcoding using nanopore MinION sequencing

**DOI:** 10.1101/2020.05.21.108852

**Authors:** Bilgenur Baloğlu, Zhewei Chen, Vasco Elbrecht, Thomas Braukmann, Shanna MacDonald, Dirk Steinke

**Author notes:** Corresponding author: Bilgenur Baloglu.

## Abstract

Metabarcoding has become a common approach to the rapid identification of the species composition in a mixed sample. The majority of studies use established short-read high-throughput sequencing platforms. The Oxford Nanopore MinION™, a portable sequencing platform, represents a low-cost alternative allowing researchers to generate sequence data in the field. However, a major drawback is the high raw read error rate that can range from 10% to 22%.

To test if the MinION™ represents a viable alternative to other sequencing platforms we used rolling circle amplification (RCA) to generate full-length consensus DNA barcodes (658bp of cytochrome oxidase I - COI) for a bulk mock sample of 50 aquatic invertebrate species. By applying two different laboratory protocols, we generated two MinION™ runs that were used to build consensus sequences. We also developed a novel Python pipeline, ASHURE, for processing, consensus building, clustering, and taxonomic assignment of the resulting reads.

We were able to show that it is possible to reduce error rates to a median accuracy of up to 99.3% for long RCA fragments (>45 barcodes). Our pipeline successfully identified all 50 species in the mock community and exhibited comparable sensitivity and accuracy to MiSeq. The use of RCA was integral for increasing consensus accuracy, but it was also the most time-consuming step during the laboratory workflow and most RCA reads were skewed towards a shorter read length range with a median RCA fragment length of up to 1262bp. Our study demonstrates that Nanopore sequencing can be used for metabarcoding but we recommend the exploration of other isothermal amplification procedures to improve consensus length.

## Introduction

DNA metabarcoding uses high-throughput sequencing (HTS) of DNA barcodes to quantify the species composition of a heterogeneous bulk sample. It has gained importance in fields such as evolutionary ecology (Lim et al. 2016), food safety (Staats et al. 2016), disease surveillance (Batovska et al. 2018), and pest identification (Sow et al. 2019). Most metabarcoding studies to date have used short-read platforms such as the Illumina MiSeq (Piper et al. 2019). New long-read instruments such as the Pacific Biosciences Sequel platform could improve taxonomic resolution (Tedersoo et al. 2017; Heeger et al. 2018) through long high-fidelity DNA barcodes. Long read nanopore devices are becoming increasingly popular because these devices are low-cost and portable (Menegon et al. 2017).

Nanopore sequencing is based on the readout of ion current changes occurring when single-stranded DNA passes through a protein pore such as alpha-hemolysin (Deamer et al. 2016). Each nucleotide restricts ion flow through the pore by a different amount, enabling base-calling via time series analysis of the voltage across a nanopore. (Clarke et al. 2009). The first commercially available instrument, Oxford Nanopore Technologies’ MinION™, is a portable, low-cost sequencing platform that can produce long reads (10 kb to 2 Mb reported; Nicholls et al. 2019). The low capital investment costs (starting at $1,000 US) have made this device increasingly popular among scientists working on molecular species identification (Parker et al. 2017, Kafetzopoulou et al. 2018, Loit et al. 2019), disease surveillance (Quick et al. 2016), and whole-genome reconstruction (Loman et al. 2015). However, a major drawback is the high raw read error rate which reportedly ranges from 10-22% (Jain et al. 2015, Sović et al. 2016, Jain et al. 2018, Kono and Arakawa, 2019, Krehenwinkel et al. 2019), a concern when investigating the within-species diversity or the diversity of closely related species.

However, with consensus sequencing strategies, nanopore instruments can also generate high fidelity reads for shorter amplicons (Simpson et al. 2017, Pomerantz et al. 2018, Rang et al. 2018). Clustering of corresponding reads is accomplished by using a priori information such as reference genomes (Vaser et al. 2017), primer indices marking each sample (Srivathsan et al. 2018), or spatially related sequence information, which can be encoded using DNA amplification protocols such as loop-mediated isothermal amplification (LAMP) (Mori & Notomi, 2009) or rolling circle amplification (RCA) (McNaughton et al. 2019). RCA is based on the circular replication of single-stranded DNA molecules. A series of such replicated sequences can be used to build consensus sequences with an accuracy of up to 99.5% (Li et al. 2016, Calus et al. 2017, Volden et al. 2018).

The combination of metabarcoding and nanopore sequencing could allow researchers to generate barcode sequence data for community samples in the field, without the need to transport or ship samples to a laboratory. So far only a small number of studies have demonstrated the suitability of MinION™ for metabarcoding using samples of very low complexity, e.g., comprising of three (Batovska et al. 2018), 6 -11 (Voorhuijzen-Harink et al. 2019), or nine species (Krehenwinkel et al. 2019).

For this study we used a modified RCA protocol (Li et al. 2016) for nanopore consensus sequencing of full-length DNA barcodes (658bp of cytochrome oxidase I - COI) from a bulk sample of 50 aquatic invertebrate species to explore the feasibility of nanopore sequencing for metabarcoding. We also developed a new Python pipeline to explore error profiles of nanopore consensus sequences, mapping accuracy, and overall community representation of a complex bulk sample.

## Methods

### Mock community preparation

We constructed a mock community of 50 freshwater invertebrate specimens collected with kick-nets in Southern Ontario and Germany. Collection details are recorded in the public dataset DS-NP50M on Barcode of Life Data Systems (BOLD, http://www.boldsystems.org, see Ratnasingham & Hebert 2007). A small piece of tissue was subsampled from each specimen (Arthropoda: a leg or a section of a leg; Annelida: a small section of the body; Mollusca: a piece of the mantle) and the DNA was extracted in 96-well plates using membrane-based protocols (Ivanova et al. 2006, Ivanova et al. 2008). The 658 bp barcode region of COI was amplified using the following thermal conditions: initial denaturation at 94°C for 2 min followed by 5 cycles of denaturation for 40 s at 94°C, annealing for 40 s at 45°C and extension for 1 min at 72°C; then 35 cycles of denaturation for 40 s at 94°C with annealing for 40 s at 51°C and extension for 1 min at 72°C; and a final extension for 5 min at 72°C (Ivanova et al. 2006). The 12.5 μl PCR reaction mixes included 6.25 μl of 10% trehalose, 2.00 μl of ultrapure water, 1.25 μl 10X PCR buffer [200 mM Tris-HCl (pH 8.4), 500 mM KCl], 0.625 μl MgCl (50 mM), 0.125 μl of each primer cocktail (0.01 mM, C_LepFolF/C_LepFolR (Hernández-Triana et al. 2014) and for Mollusca C_GasF1_t1/GasR1_t1 (Steinke et al. 2016)), 0.062 μl of each dNTP (10 mM), 0.060 μl of Platinum® Taq Polymerase (Invitrogen), and 2.0 μl of DNA template. PCR amplicons were visualized on a 1.2% agarose gel E-Gel^®^ (Invitrogen) and bidirectionally sequenced using sequencing primers M13F or M13R and the BigDye^®^Terminator v.3.1 Cycle Sequencing Kit (Applied Biosystems, Inc.) on an ABI 3730xl capillary sequencer following manufacturer’s instructions. Bi-directional sequences were assembled and edited using Geneious 11 (Biomatters). For specimens without a species-level identification, we employed the Barcode Index Number (BIN) system that assigns each specimen to a species proxy using the patterns of sequence variation at COI (Ratnasingham & Hebert, 2013). With this approach, we selected a total of 50 OTUs with 15% or more K2P COI distance (Kimura, 1980) from other sequences for the mock sample. A complete list of specimens, including taxonomy, collection details, sequences, BOLD accession numbers, and Nearest Neighbour distances are provided in Supplementary Table S1.

### Bulk DNA extraction

The remaining tissue of the mock community specimens was dried overnight, pooled, and subsequently placed in sterile 20mL tubes containing 10 steel beads (5mm diameter) to be homogenized by grinding at 4000 rpm for 30-90 min in an IKA ULTRA TURRAX Tube Drive Control System (IKA Works, Burlington, ON, Canada). A total of 22.1 mg of homogenized tissue was used for DNA extraction with the Qiagen DNeasy Blood and Tissue kit (Qiagen, Toronto, ON, Canada) following the manufacturer’s instructions. DNA extraction success was verified on a 1% agarose gel (100 V, 30 min) and DNA concentration was quantified using the Qubit HS DNA Kit (Thermo Fisher Scientific, Burlington, ON, Canada).

### Metabarcoding using Illumina Sequencing

For reference, we used a common metabarcoding approach with a fusion primer-based two-step PCR protocol (Elbrecht & Steinke 2019). During the first PCR step, a 421 bp region of the Cytochrome c oxidase subunit I (COI) was amplified using the BF2/BR2 primer set (Elbrecht & Leese 2017). PCR reactions were carried out in a 25 µL reaction volume, with 0.5 µL DNA, 0.2 µM of each primer, 12.5 µL PCR Multiplex Plus buffer (Qiagen, Hilden, Germany). The PCR was carried out in a Veriti thermocycler (Thermo Fisher Scientific, MA, USA) using the following cycling conditions: initial denaturation at 95 °C for 5 min; 25 cycles of: 30 sec at 95 °C, 30 sec at 50 °C and 50 sec at 72 °C; and a final extension of 5 min at 72 °C. One µL of PCR product was used as the template for the second PCR, where Illumina sequencing adapters were added using individually tagged fusion primers (Elbrecht & Steinke 2019). For the second PCR, the reaction volume was increased to 35 µL, the cycle number reduced to 20, and extension times increased to 2 minutes per cycle. PCR products were purified and normalized using SequalPrep Normalization Plates (Thermo Fisher Scientific, MA, USA, Harris et al. 2010) according to manufacturer protocols. Ten µL of each normalized sample was pooled, and the final library cleaned using left-sided size selection with 0.76x SPRIselect (Beckman Coulter, CA, USA). Sequencing was carried out by the Advances Analysis Facility at the University of Guelph using a 600 cycle Illumina MiSeq Reagent Kit v3 and 5% PhiX spike in. The forward read was sequenced for an additional 16 cycles (316 bp read).

The resulting sequence data were processed using the JAMP pipeline v0.67 (github.com/VascoElbrecht/JAMP). Sequences were demultiplexed, paired-end reads merged using Usearch v11.0.667 with fastq_pctid=75 (Edgar 2010), reads below the read length threshold (414bp) were filtered and primer sequences trimmed both by using Cutadapt v1.18 with default settings (Martin 2011). Sequences with poor quality were removed using an expected error value of 1 (Edgar & Flyvbjerg 2015) as implemented in Usearch. MiSeq reads, including singletons, were clustered using cd-hit-est (Li & Godzik, 2006) with parameters: -b 100 -c 0.95 -n 10. Clusters were subsequently mapped against the mock community data as well as against the BOLD COI reference library.

### Metabarcoding using Nanopore sequencing

We used a modified intramolecular-ligated Nanopore Consensus Sequencing (INC-Seq) approach (Li et al. 2016) that employs rolling circle amplification (RCA) of circularized templates to generate linear tandem copies of the template to be sequenced on the nanopore platform. An initial PCR was prepared in 50μl reaction volume with 25μl 2× Multiplex PCR Master Mix Plus (Qiagen, Hilden, Germany), 10pmol of each primer (for 658 bp COI barcode fragment – Supplementary Table S2), 19μl molecular grade water and 4μl DNA. We used a Veriti thermocycler (Thermo Fisher Scientific, MA, USA) and the following cycling conditions: initial denaturation at 98°C for 30 secs, 35 cycles of (98°C for 30 secs, 59°C for 30 secs, 72°C for 30 secs), and a final extension at 72°C for 2 min. Amplicons were purified using SpriSelect (Beckman Coulter, CA, USA) with a sample to volume ratio of 0.6x and quantified. Purified amplicons were self-ligated to form plasmid like structures using Blunt/TA Ligase Master Mix (NEB, Whitby, ON, Canada) following manufacturer’s instructions. Products were subsequently treated with the Plasmid-SafeTM ATP-dependent DNAse kit (Lucigen Corp, Middleton, WI, USA) to remove remaining linear molecules. Final products were again purified with SpriSelect at a 0.6x ratio and quantified using the High Sensitivity dsDNA Kit on a Qubit fluorometer (Thermo Fisher Scientific, MA, USA). Rolling Circle Amplification (RCA) was performed for six 2.5 μL aliquots of circularized DNA plus negative controls (water) using the TruePrime™ RCA kit (Expedeon Corp, San Diego, CA, USA) following manufacturer’s instructions. After initial denaturation at 95°C for three minutes, RCA products were incubated for 2.5 to 6 hours at 30°C. The DNA concentration was measured after every hour. RCA was stopped once 60-70 ng/ul of double-stranded DNA was reached. Subsequently, RCA products were incubated for 10 min at 65°C to inactivate the enzyme. We performed two experiments under varying RCA conditions (Protocol A and B, detailed in Table 1), such as RCA duration (influences number of RCA fragments), fragmentation duration, and fragmentation methods.

**Table 1:** Varying RCA conditions for experimental protocols A and B

| Dataset | Protocol A | Protocol B |
| --- | --- | --- |
| RCA duration (hrs) | 5 | 6 |
| Number of target sequences per RCA fragment | 12 | 15 |
| Enzymatic branching (min) | 5 | 2 |
| Mechanical fragmentation | 4200 rpm, 2 min | None |
| Primer pairs used | HCOA-LCO, HCOC2-LCOC2 | HCOA2-LCOA2, HCOC2-LCOC2 |

Protocol A followed Li et al. (2016) by incubating 65μL of pooled RCA product with 2μL (20 units) of T7 Endonuclease I (NEB, M0302S, VWR Canada, Mississauga, ON, Canada) at room temperature for 10 min of enzymatic debranching, followed by mechanical shearing using a Covaris g-TUBETM (D-Mark Biosciences, Toronto, ON, Canada) at 4200 rpm for 1 min on each side of the tube or until the entire reaction mix passed through the fragmentation hole. Protocol B is a more modified approach to counteract the overaccumulation of smaller DNA fragments. Here we did only 2 min of enzymatic debranching with no subsequent mechanical fragmentation. To verify the size of fragments after shearing, sheared products for both protocols were run on a 1% agarose gel at 100 V for 1 hour. DNA damage was repaired by incubating 53.5μL of the product with 6.5μL of FFPE DNA Repair Buffer and 2μL of NEBNext FFPE Repair mix (VWR Canada, Mississauga, ON, Canada) at 20°C for 15. The final product was purified using SpriSelect at a 0.45x ratio and quantified using a Qubit fluorometer.

For sequencing library preparation, we used the Nanopore Genomic Sequencing Kit SQK-LSK308 (Oxford Nanopore, UK). First, the NEBNext Ultra II End Repair/dA Tailing kit (NEB, Whitby, ON, Canada) was used to end repair 1000 ng of sheared genomic DNA (1 microgram of DNA in 50μl nuclease-free water, 7μl of Ultra II End-Prep Buffer, 3μl Ultra II End-Prep Enzyme Mix in a total volume of 60μl). The reaction was incubated at 20°C for 5 min and heat-inactivated at 65°C for another 5 min. Resulting DNA was purified using SpriSelect at a 1:1 ratio according to the SQK-LSK308 protocol. Then it was eluted in 25μl of nuclease-free water and quantified with a recovery aim of >70 ng/μl. Blunt/TA Ligase Master Mix (NEB, Whitby, ON, Canada) was used to ligate native barcode adapters to 22.5μl of 500 ng end-prepared DNA at room temperature (10 min). DNA was purified using a 1:1 volume of SpriSelect beads and eluted in 46μl nuclease-free water before the second adapter ligation. For each step, the DNA concentration was measured. The library was purified with ABB buffer provided in the SQK□LSK308 kit (Oxford Nanopore, Oxford Science Park, UK). The final library was then loaded onto a MinION flow cell FLO-MIN107.1 (R9.5) and sequenced using the corresponding workflow on MinKNOW(tm). Base-calling was performed using Guppy 3.2.2 in CPU mode with the dna_r9.5_450bps_1d2_raw.cfg model.

We designed a new Python (v3.7.6) pipeline, termed ASHURE (A safe heuristic under Random Events) to process RCA reads and to build consensus sequences (Suppl Fig 1). Detailed information is available on GitHub: https://github.com/BBaloglu/ASHURE. The pipeline uses the OPTICS algorithm (Ankerst et al. 1999) for clustering and t-distributed stochastic neighbor embedding (Maaten & Hinton, 2014) for dimensionality reduction and visualization. Sequence alignments were conducted using minimap2 (Li, 2018) and SPOA (Vaser et al. 2017). Correlation coefficients were determined through ASHURE using both the Numpy (van der Walt et al. 2011) and the Pandas package (McKinney 2010). The Pipeline also includes comparisons of consensus error to several parameters, such as RCA length, UMI error, and cluster center error as well as accuracy determination. The error was calculated by dividing edit distance to the length of the shorter sequence that was compared.

We also calculated median accuracy and number of detected species using the R2C2 (Rolling Circle Amplification to Concatemeric Consensus) post-processing pipeline C3POa (Concatemeric Consensus Caller using partial order alignments) for consensus calling (Volden et al. 2018). C3POa generates two kinds of output reads: 1) Consensus reads if the raw read is sufficiently long to cover an insert sequence more than once and 2) Regular “1D” reads if no splint sequence could be detected in the raw read (Adams et al. 2019). We only used consensus reads for downstream analysis. Unlike ASHURE, C3POa does not report information on the RCA fragment length, hence we were not able to make direct comparisons for different thresholds.

## Results

### Mock community

Many collected specimens could not be readily identified to species level. Consequently, we employed the Barcode Index Number (BIN) system which examines patterns of sequence variation at COI to assign each specimen to a species proxy (Ratnasingham & Hebert, 2013). We retrieved 50 BINs showing >15% COI sequence divergence from their nearest neighbor under the Kimura 2□parameter model (Kimura, 1980). The resulting freshwater macrozoobenthos mock community included representatives of 3 phyla, 12 orders, and 27 families. COI sequences have been deposited on NCBI Genbank under the Accession Numbers MT324068-MT324117. Further specimen details can be found in the public dataset DS-NP50M (<u>dx.doi.org/10.5883/DS-NP50M</u>) on BOLD.

### Metabarcoding using Illumina Sequencing

All samples showed good DNA quality. Illumina MiSeq sequencing generated an average of 204 797 paired-end reads per primer combination. Raw sequence data are available under the NCBI SRA accession number SRR9207930. We recovered 49 of 50 OTUs present in our mock community (Fig. 1D). We obtained a total of 845 OTUs (OTU table including sequences, read counts, and assigned taxonomy is available as Supplementary Table S3) mostly contaminants that were in part also obtained with nanopore sequencing.

**Figure 1:**
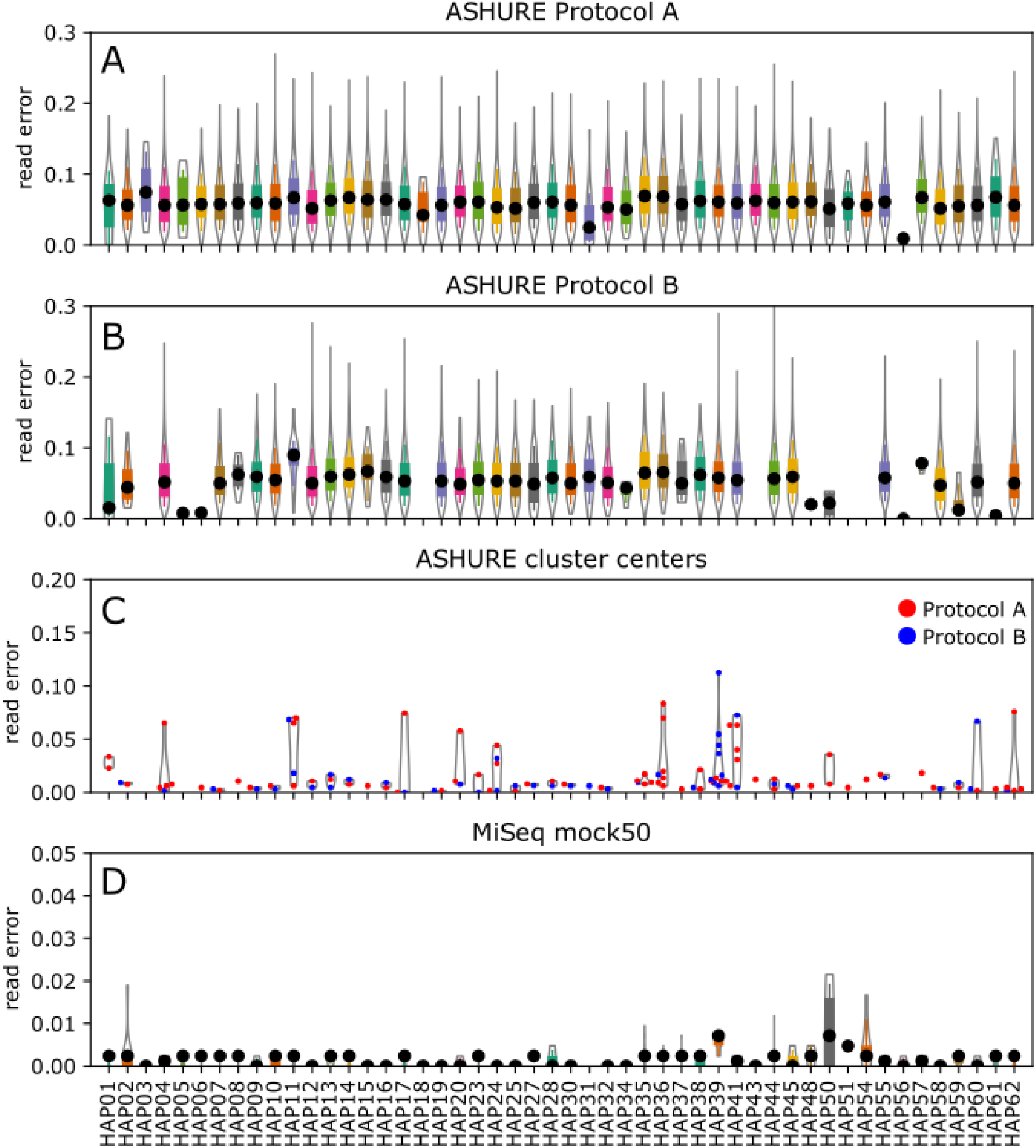
Nanopore sequencing read error per species for (A) Protocol A and (B) Protocol B obtained with ASHURE using all reads. (C) Nanopore sequencing read error obtained with OPTICS in ASHURE using cluster centers for each RCA condition. (D) MiSeq sequencing read error per species.

### Metabarcoding using Nanopore sequencing

Nanopore sequencing with the MinION delivered 746,153/2,756 and 499,453/1,874 1D/1D^2^ reads for Protocols A and B (SRA PRJNA627498), respectively. The 1D approach only sequences one template DNA strand, whereas with the 1D^2^ method both complementary strands are sequenced, and the combined information is used to create a higher quality consensus read (Cornelis et al. 2019). Because of the low read output for 1D^2^ reads, our analyses focused on 1D data. Most reads were skewed towards a shorter read length range (Figure 2) with a median RCA fragment length of 1262bp for Protocol A and 908 bp for Protocol B.

**Figure 2:**
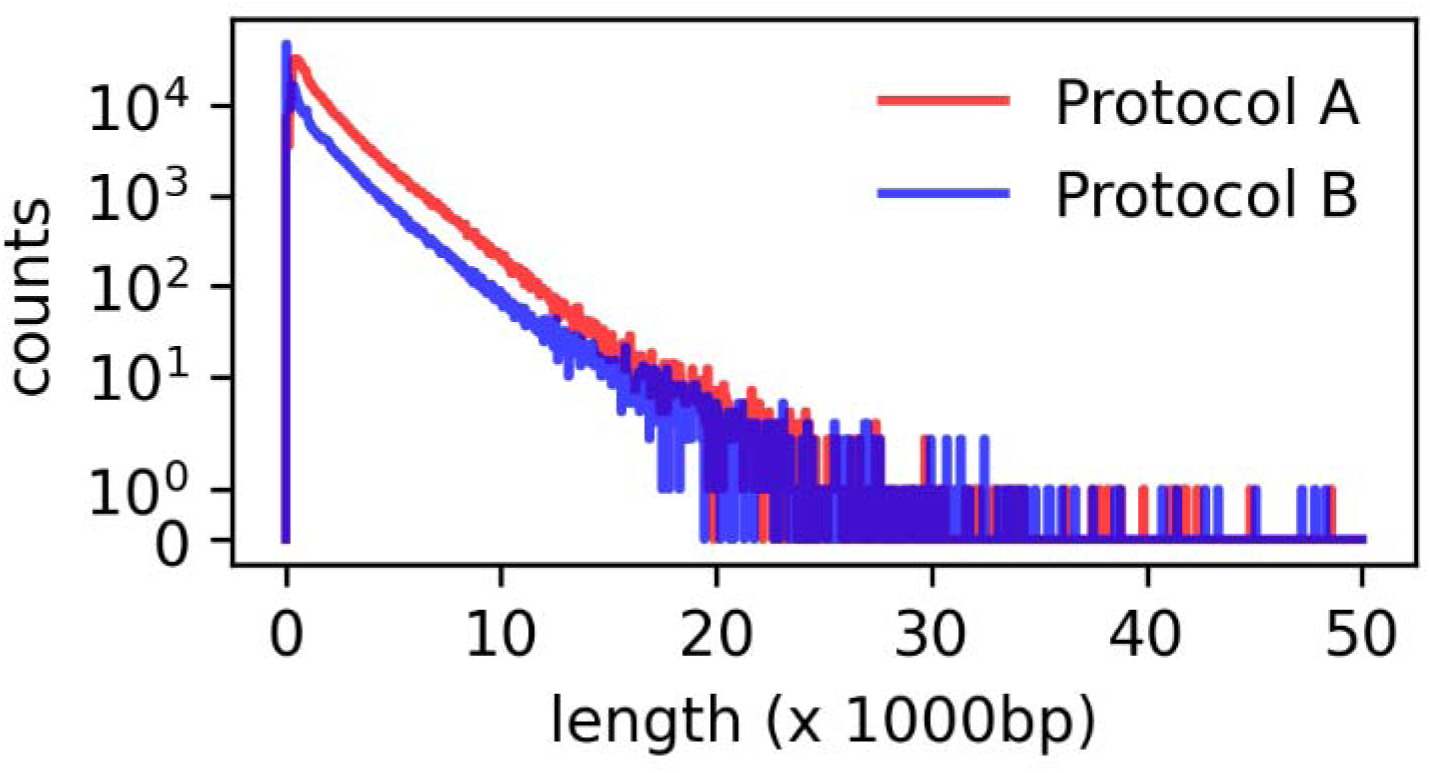
Read length distribution for both sequencing protocols. The number of reads is provided in a logarithmic scale on the y-axis.

With flexible filtering (number of targets per RCA fragment = 1 or more), ASHURE results provided a median accuracy of 92.16% for Protocol A and 92.87% for Protocol B (see Table 2, Figures 1A-B). Using ASHURE, we observed a negative, non-significant correlation between consensus median error and the number of RCA fragments (Pearson’s r for Protocol A: -0.247, Protocol B: -0.225). For both protocols, we found a positive, non-significant correlation between consensus median error and primer error (Pearson’s r for Protocol A: 0.228, Protocol B: 0.375) and between consensus median error and cluster center error (see Figures 3B-C; Pearson’s r for Protocol A: 0.770, Protocol B: 0.274). We obtained median accuracy values of >95% for 1/5^th^ of the OTUs in Protocol A and half of the OTUs in Protocol B for flexible filtering. Increasing the number of RCA fragments to 15 or more came with the trade-off of detecting fewer OTUs (from 50 to 36 for Protocol A and 50 to 38 for Protocol B). At the same time, median accuracy values increased to 97.4% and 97.6% for Protocol A and B, respectively. With more stringent filtering (number of targets per RCA fragment = 45 or more), median accuracy improved up to 99.3% for both Protocol A and B but with the trade-off of an overall reduced read output and a reduced number of species recovered (Table 2).

**Table 2:** Consensus reads, median accuracy, and the number of OTUs/species detected at different thresholds for Protocol A and B analyzed with ASHURE and C3POa.

| ASHURE pipeline |  |  |  |  |  |  |
| --- | --- | --- | --- | --- | --- | --- |
|  | Protocol A |  |  | Protocol B |  |  |
| Consensus read criterium | # of reads | Median accuracy (%) | # of OTUs detected | # of reads | Median accuracy (%) | # of OTUs detected |
| unfiltered | 269,620 | 93.6 | 198 | 245,827 | 93.4 | 188 |
| post filtering non-target data based on MiSeq | 199,709 | 92.16 | 50 | 214,782 | 92.87 | 50 |
| RCA > 15 | 1,434 | 97.39 | 36 | 2,884 | 97.62 | 38 |
| RCA > 20 | 292 | 97.86 | 28 | 1,009 | 98.10 | 34 |
| RCA > 25 | 78 | 98.22 | 19 | 455 | 98.35 | 30 |
| RCA > 30 | 20 | 98.46 | 11 | 217 | 98.57 | 26 |
| RCA > 35 | 7 | 99.05 | 5 | 106 | 98.82 | 22 |
| RCA > 40 | 3 | 99.52 | 2 | 57 | 99.05 | 18 |
| RCA > 45 | 2 | 99.60 | 2 | 30 | 99.29 | 13 |
| RCA > 50 | 1 | 99.68 | 1 | 21 | 98.82 | 8 |
| C3POa |  |  |  |  |  |  |
| unfiltered | 322,884 | 94.5 | 180 | 128,353 | 94.7 | 118 |
| post filtering non-target data based on MiSeq | 314,041 | 94.5 | 50 | 124,131 | 94.7 | 40 |

**Figure 3:**
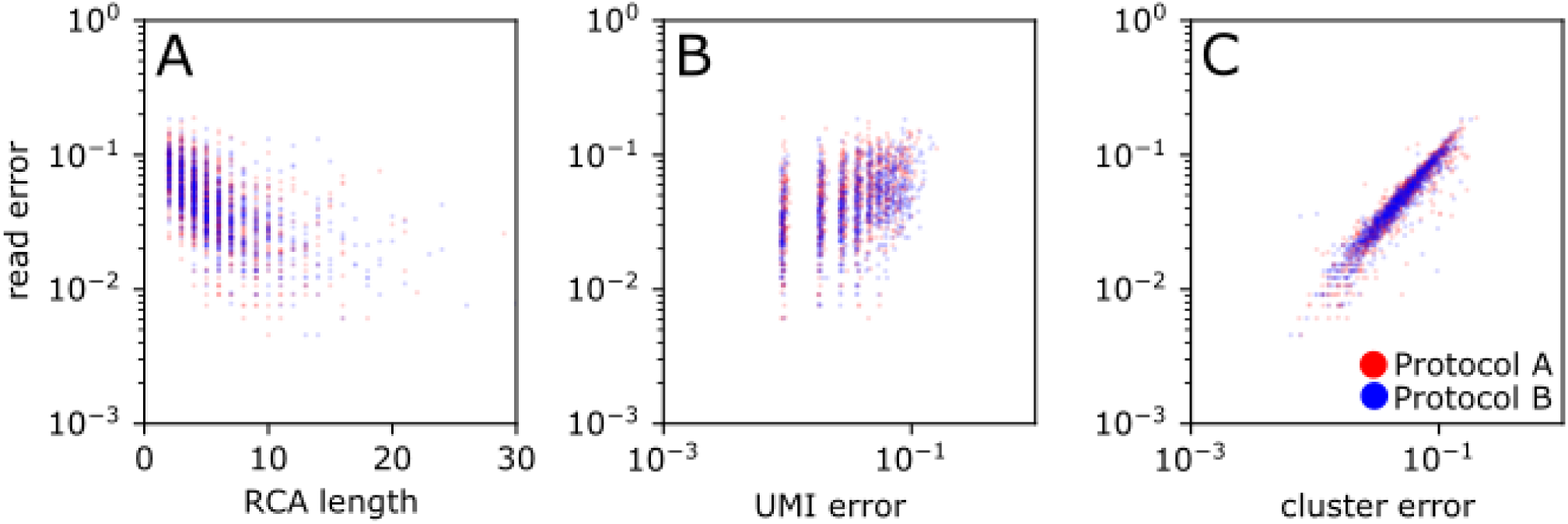
Comparison of consensus error versus (A) RCA length, (B) UMI error, and (C) cluster center error using the ASHURE pipeline for two RCA conditions.

We mapped the 845 OTUs found in the MiSeq dataset to the Nanopore reads and removed contaminants, (69,911 for Protocol A and 31,045 reads for Protocol B) using ASHURE. With Miseq, we were able to detect 49 out of 50 of the mock species, whereas all 50 mock community species were detected in both nanopore sequencing protocols A and B. Using the MiSeq dataset, we also removed contaminants from the consensus reads obtained with C3POa (8,843 for Protocol A and 4,222 reads for Protocol B). Using C3POa, we retained a lower number of consensus reads than with ASHURE for Protocol B (see Table 2), but the median consensus accuracy using flexible filtering was similar (94.5-94.7% Protocol A and B). The median accuracy when including all consensus reads was higher for C3POa than ASHURE in both Protocol A and B. Overall the two pipelines showed similar performance in consensus read error profile (Supplementary Figures 2A-D, Supplementary Figure 3). As for Protocol B, ASHURE detected a higher number of mock community species (see Table 2).

The read error of all consensus reads (Figures 1A-B) spanned a wide range (0-10% error). Running OPTICS, a density-based clustering algorithm, on the consensus reads enabled us to identify cluster centers (Fig. 1C), which possessed comparable accuracy to MiSeq (Fig. 1D). Figures 3A-C show comparisons of consensus error with RCA length, UMI error, and cluster center error. We found that cluster center error correlated better with consensus error, particularly for Protocol A (Pearson’s r: 0.770), (see Figure 3C). To visualize why OPTICS can identify high fidelity cluster centers, five OTUs were randomly selected and clustered at different RCA fragment lengths (Figure 4). T-distributed stochastic neighbor embedding (t-SNE) was used to visualize the co-similarity relationship of this collection of sequences in two dimensions (Figures 4B-F). Closely related sequences clustered together and corresponded to the OTUs obtained by OPTICS. Clustering of raw reads resulted in less informative clusters, where OTUs were not well separated and cluster membership did not match that of the true species (Fig. 4C). The clustering of reads with increasing RCA length cut-off resulted in clusters that had more distinct boundaries (Figures 4D-F). These clusters corresponded to the true haplotype sequences (Fig. 4F) and contained the de novo cluster centers and true OTU sequences at their centroids. The OPTICS algorithm successfully extracted the OTU structure embedded in a co-similarity matrix, flagged low fidelity reads that were in the periphery of each cluster, and ordered high fidelity reads to the center of the clusters (Fig. 4B).

**Figure 4:**
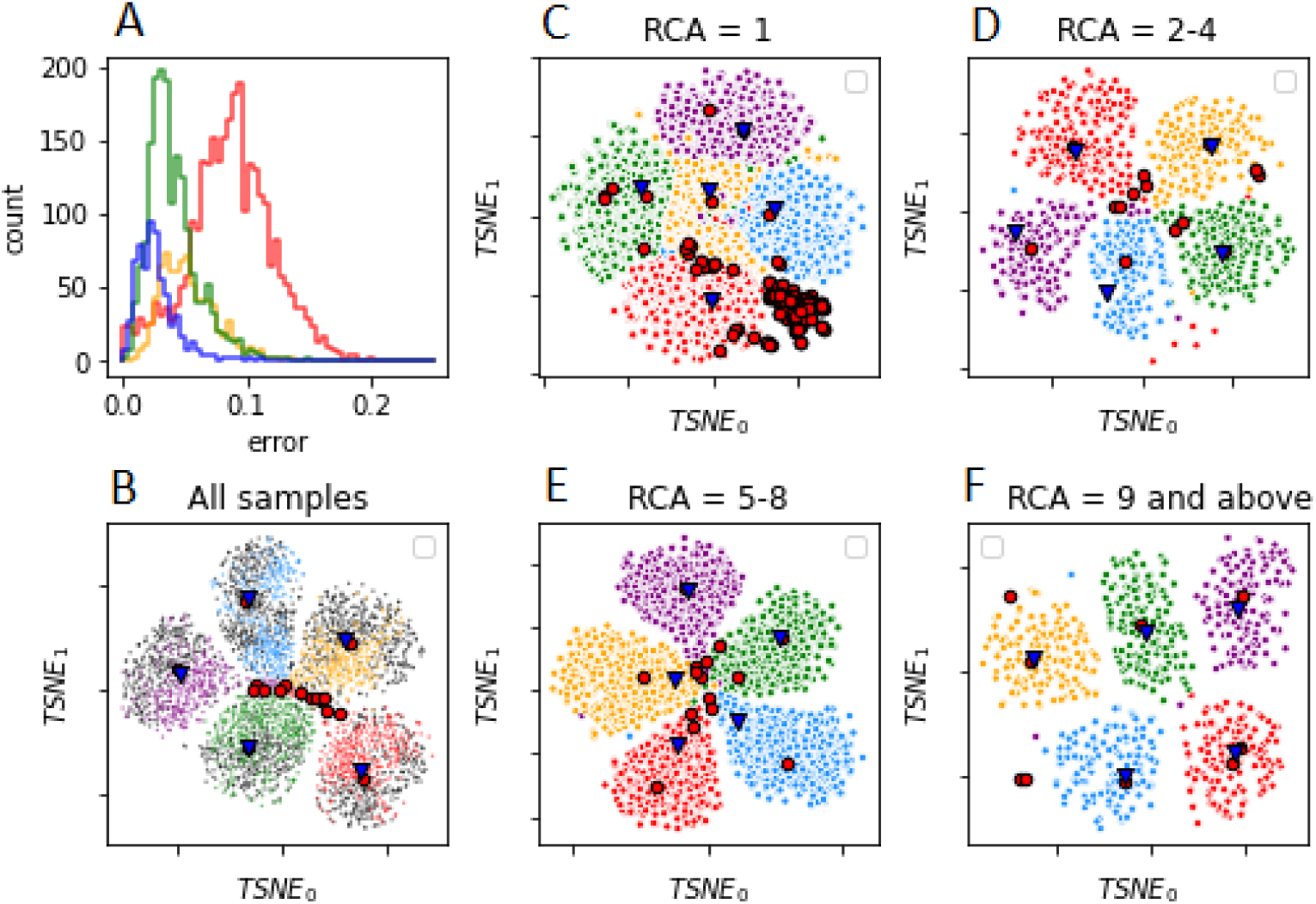
tSNE visualization of reference-free clustering using OPTICS for five randomly selected haplotypes. (A) The number of reads and percentage of error for each filtering criteria, red: reads with 1 RCA fragment, yellow: reads with 2-4 RCA fragments, green: reads with 5-8 RCA fragments, and blue: reads with 9 or more RCA fragments. tSNE visualization of OPTICS clusters for reads with (B) no filtering, (C) one RCA fragment, (D) 2-4 RCA fragments, (E) 5-8 RCA fragments, (F) 9 and more RCA fragments. True haplotypes (blue triangles) and cluster centers obtained with reference-free clustering (red circles) overlap more as the number of RCA fragments increases. Colors in B-F correspond to: HAP04 (red), HAP11 (blue), HAP17 (purple), HAP39 (orange), HAP41 (green). Grey dots in (B) indicate outliers.

## Discussion

This study introduces a workflow for DNA metabarcoding of freshwater organisms using the Nanopore MinION™ sequencing platform. We were able to show that it is possible to mitigate the high error rates associated with nanopore-based long-read single-molecule sequencing by using rolling circle amplification with a subsequent assembly of consensus sequences leading to a median accuracy of up to 99.3% for long RCA fragments (>45 barcodes).

We were able to retrieve all OTUs of the mock community assembled for this study. Our mock sample species had at least 15% genetic distance to each other and with ASHURE we were able to retrieve them both under relaxed and strict filtering conditions. This will likely change if a sample includes species that are more closely related with average distances of 2-3%. Although both of our experimental protocols were successful, we observed a higher number of consensus reads, detected species overall and median accuracy for Protocol B which used a higher number of RCA replicates as input DNA, had no mechanical fragmentation step, and a reduced duration of enzymatic debranching (Table 2). We recommend adopting our Protocol B workflow and using strict filtering in the ASHURE pipeline, e.g. a minimum of 15 barcodes per RCA fragment. We used the Illumina MiSeq platform to identify by-products or contaminants as well as for comparison with nanopore sequencing. In terms of accuracy the MiSeq platform performs slightly better (Figure 1C and D). However, the improved error rates clearly make the MinION™ a more cost-effective and mobile alternative.

Consensus sequence building is the critical step for achieving high accuracy with MinION™ reads. Raw outputs of Nanopore sequencing are improving (Volden et al. 2018) and as read accuracy further improves, so will the quality of consensus sequences. We show that RCA is integral for increasing consensus accuracy, but it is also the most time-consuming step during the laboratory workflow, e.g. with 60-70 ng/ul of input DNA 5-6 hours of RCA were necessary to achieve reasonable results. Our results display a trade-off between median consensus accuracy and the detection of species, particularly due to not having enough long reads (see Table 2, Fig. 2). However, despite most reads being relatively short, we observed an inverse correlation between RCA length and the consensus error rate (Fig. 3A). For further improvement of consensus sequence accuracy, the proportion of longer reads needs to be maximized. For more time-sensitive studies on metabarcoding with Nanopore sequencing, e.g. field-based studies, we suggest modifying the RCA duration based on the complexity of the sample. However, given some of the RCA weaknesses, we recommend the exploration of other isothermal amplification procedures such as LAMP (Imai et al. 2017), multiple displacement amplification, (MDA) (Hansen et al. 2018), or recombinase polymerase amplification, (RPA) (Donoso & Valenzuela, 2018).

Previous studies using circular consensus approaches to Nanopore sequencing, such as INC-seq (Li et al. 2016) and R2C2 (Volden et al. 2018) have already shown improvements in read accuracy. We compared our pipeline ASHURE with C3POa, the post-processing pipeline for R2C2 with a reported median accuracy of 94% (Volden et al. 2018). C3POa data processing includes the detection of DNA splint sequences and the removal of short (<1,000 kb) and low-quality (Q < 9) reads (Volden et al. 2018). With C3POa, a raw read is only used for consensus calling if one or more specifically designed splint sequences are detected within it (Volden et al. 2018). Instead of splint sequences we used primer sequences to identify reads for further consensus assembly. Both C3POa and ASHURE showed similar accuracy for our datasets, but C3POa detected fewer species in our Protocol B experiment. Using ASHURE, we were only able to detect 43.4% and 7% of the reads with both primers attached in Protocol A and B, respectively. This points to some issues with the RCA approach and might explain why C3POa generated fewer numbers of consensus reads in Protocol B, as the number of detected sequences was very low. Initially we assumed that increasing the unique molecular identifier (UMI) length for our primers would be useful not only for consensus calling but also for identifying, quantifying, and filtering erroneous consensus reads. However, within the small percentage of reads with both primers attached, we did not find a strong correlation between the UMI error and the consensus read error (Figure 3B).

Several MinION™ studies have implemented a reference-free approach for consensus calling, however, these studies are limited to tagged amplicon sequencing that allows for sequence-to-specimen association (Srivathsan et al. 2018, Calus et al. 2018; Pomerantz et al. 2018; Srivathsan et al. 2019). Such an approach can be useful for species-level taxonomic assignment (Benítez-Páez et al. 2016) and even species discovery (Srivathsan et al. 2019). Our pipeline uses density-based clustering which is a promising approach when studying species diversity in mixed samples, particularly with Nanopore sequencing. The density-based clustering of Nanopore reads allows for a reference-free approach by grouping reads with their replicates without having to map to a reference database (Faucon et al. 2017). Conventional OTU threshold clustering approaches have shown to be a challenge for nanopore data. Either each sequence was assigned to a unique OTU, or OTU assignment failed due to the variable error profile (Ma et al. 2017), or the optimal threshold depended on the relative abundance of species in a given sample (Mafune et al. 2017). Density-based clustering is advantageous because it can adaptively call cluster boundaries based on other objects in the neighborhood (Ankerst et al. 1999). Clusters correspond to the regions in which the objects are dense, and the noise is regarded as the regions of low object density (Ankerst et al. 1999). For DNA sequences, such a clustering approach requires sufficient read coverage around a true amplicon so that the novel clusters can be detected and are not treated as noise. With sufficient sample size, density-based approaches can allow us to obtain any possible known or novel species clusters with high accuracy and without the need for a reference database. ASHURE is not limited to RCA data, as it performs a search for primers in the sequence data, splits the reads at primer binding sites, and stores the information on start and stop location of the fragment as well as its orientation. The pipeline can be used to process outputs of other isothermal amplification methods generating concatenated molecules by simply providing primer/UMI sequences that link each repeating segment.

## Conclusion

This study demonstrates the feasibility of bulk sample metabarcoding with Oxford Nanopore sequencing using a modified molecular and novel bioinformatics workflow. We highly recommend the use of isothermal amplification techniques to obtain longer repetitive reads from a bulk sample. With our pipeline ASHURE, it is possible to obtain high-quality consensus sequences with up to 99.3% median accuracy and to apply a reference-database free approach using density-based clustering. This study was based on aquatic invertebrates, but the pipeline can be extended to many other taxa and ecological applications. By offering portable, highly accurate, and species-level metabarcoding, Nanopore sequencing presents a promising and flexible alternative for future bioassessment programs and it appears that we have reached a point where highly accurate and potentially field-based DNA metabarcoding with this instrument is possible.

## Acknowledgments

We thank all staff at the CBG who helped to collect the samples employed to assemble the mock community. We also would like to thank Florian Leese, Arne Beermann, Cristina Hartmann-Fatu, and Marie Gutgesell for collecting and providing specimens. This study was supported by funding through the Canada First Research Excellence Fund. The funders had no role in study design, data collection, and analysis, decision to publish, or preparation of the manuscript.

This work represents a contribution to the University of Guelph Food From Thought research program.

## Author contributions

BB, VE, TB, and DS designed the experiments; BB and SM assembled the mock community, BB did lab work; VE did the MiSeq experiment, BB and ZC analyzed the data; BB and DS wrote the manuscript, all authors contributed to the manuscript.

