## Supplementary Figures for "A workflow for accurate metabarcoding using nanopore MinION sequencing"

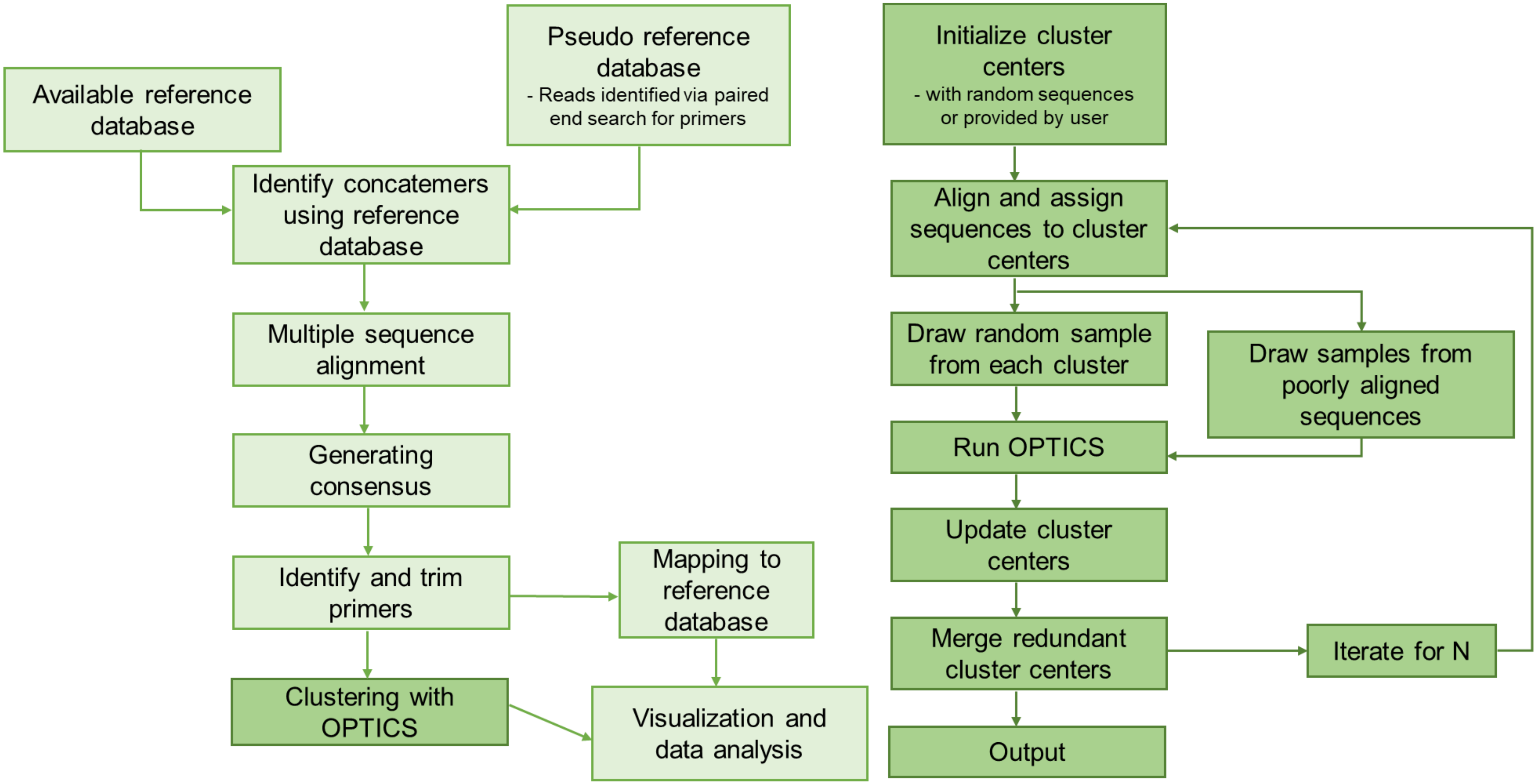


**Supplementary Figure S1:** Overview of the ASHURE bioinformatics pipeline. Clustering step is coloured in darker green and depicted in more detail on the left.

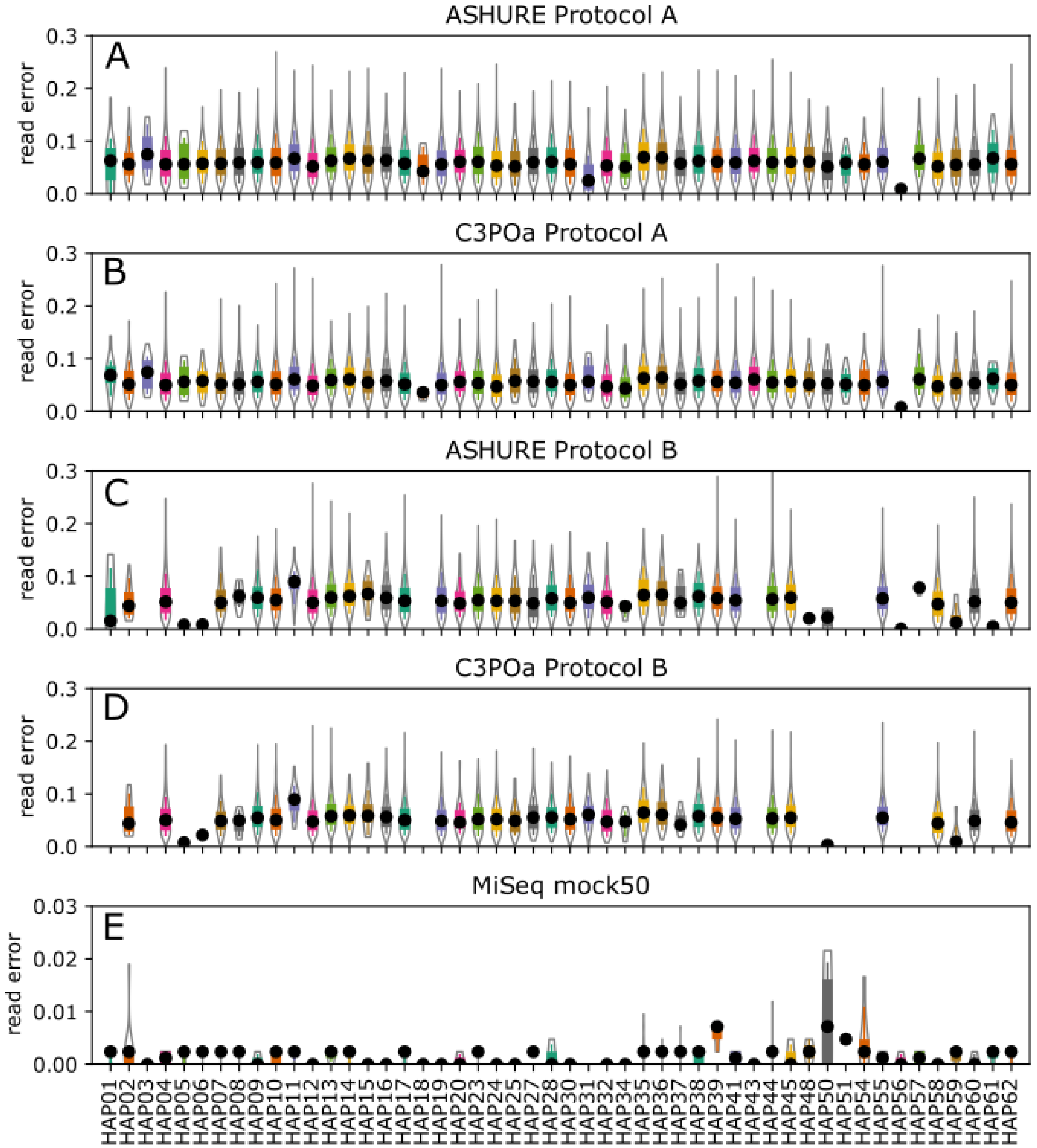


**Suppl Figure 2:** Nanopore sequencing consensus read error per species for Protocol A obtained with (A) ASHURE and (B) C3POa, for Protocol B obtained with (C) ASHURE and (D) C3POa. (E) shows sequence read error per species obtained with the Illumina MiSeq platform.

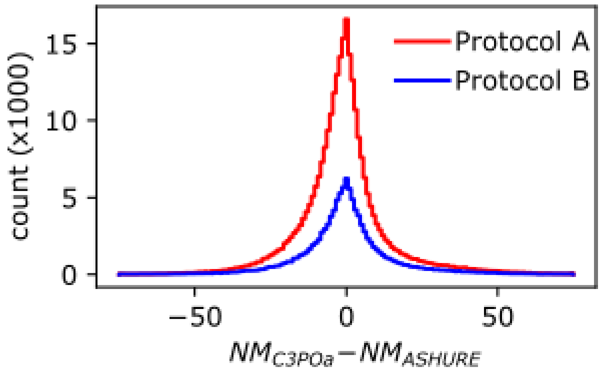


**Suppl Figure 3:** The change in the error (the number of nucleotides shown as *NM* that differ between consensus and haplotype sequence) for sequences that overlap between C3POa and ASHURE. Both pipelines perform similarly as the peak is around 0 for both protocols A and B.
